# Manatee genomics supports a special conservation area in the Guianas coastline under the influence of the Amazon River plume

**DOI:** 10.1101/552919

**Authors:** Sibelle T. Vilaça, Camilla S. Lima, Camila J. Mazzoni, Fabricio R. Santos, Benoit de Thoisy

## Abstract

The West Indian manatee (*Trichechus manatus*) occurs along the Atlantic coastline and adjacent freshwater systems of South, Central and North America, from Alagoas (Brazil) to Florida (USA) and Greater Antilles. The Amazonian manatee (*Trichechus inunguis*) is the only Sirenian adapted exclusively to freshwater and endemic to the Amazon River basin. Previous studies have reported hybrids between *T. inunguis* and *T. manatus* close to the mouth of the Amazon River, composing a likely extensive hybrid zone along the Guianas coast of South America under the influence of the Amazon River plume. We have used ddRAD SNP data, and sequences of nuclear and mtDNA loci to characterize the genomic composition of manatees along the French Guiana coastline. We found this population to be formed by introgressed or later generation interspecific hybrids, and also describe the first *T. inunguis* found outside the Amazon River basin. Our results indicate that *T. inunguis* can survive in the Amazon River plume and have colonized independent water streams of the Guianas coastline where they likely hybridize with *T. manatus*. This hypothesis offers a plausible explanation for the known extension of the hybrid zone between the two species along the Guianas coastline. It also reinforces the importance of the Amazon plume, which flows westwards to the Guianas coastline and favors the dispersion of the freshwater species. This habitat functions as a large estuary-like system that provides an ecological continuum from Amazon River mouth to the disconnected waterflows of the Guianas, which deserves a status of special conservation area.

## Introduction

Manatees (*Trichechus spp.*) are charismatic and vulnerable aquatic species that can be found in the Americas (*T. manatus* and *T. inunguis*) and coastal regions of Africa (*T. senegalensis*) (Deutsch, Self-Sullivan, & Mignucci-Giannoni, 2008). The West Indian manatee, *T. manatus*, can live in both salt and freshwater environments. With exception of the population inhabiting the northeastern coast of Brazil (Santos et al., 2016), it is also found in rivers and lake systems along its coastal distribution. Previous observations have recorded *T. manatus* up to 1100 km upstream in the Orinoco River (Castelblanco-Martinez et al., 2003) and up to 650 km in the Magdalena River in Colombia (Montoya-Ospina et al., 2001). Conversely, the Amazonian manatee (*T. inunguis*) is highly specialized to freshwater environments and has only been reported in the Amazon River basin so far.

French Guiana (FG), an overseas department of France in South America, has a stable manatee population, allegedly belonging to the *T. manatus* species, estimated to have between 50 and 250 mature individuals (UICN France et al., 2017)□. The main threats to the FG manatee population are poaching, bycatches and degradation of coastal water quality (UICN France et al., 2017). Despite a significant population decrease during the last century, recolonization of some areas has been observed (de Thoisy et al., 2003). Due to small population sizes, few catches per year are enough to maintain its declining trend and suggest a regional threatened status (UICN France et al., 2017). Sighting records are widespread in FG, both in dense and well-preserved mangroves and on the lower parts of all river systems. Important seasonality in habitat preference has been noticed in relation to strong seasonal fluctuations in the relative contributions of marine and continental waters to the estuarine waters (Castelblanco-Martínez, dos Reis, & de Thoisy, 2017). However, FG manatees also make use of large freshwater streams, with records up to 100 km inland (UICN France et al., 2017).

Previous studies involving *T. manatus* populations (Garcia-Rodriguez et al., 1998; Vianna et al., 2006; Santos et al., 2016) have identified three deeply divergent mitochondrial DNA (mtDNA) clusters composing two Evolutionary Significant Units (ESUs) that occupy two major geographic regions: Caribbean coast of Antilles, South and Central America, Mexico Gulf coast and Florida (USA) (clusters I and II); and the northeastern Atlantic coast of South America (Brazil and the Guianas), where cluster III was exclusively found (Vianna et al., 2006). However, seven manatees found along the Guianas coastline (corresponding to Guyana, French Guiana and the state of Amapá, Brazil) and one manatee found in the Amazon River close to the mouth were reported to be hybrids with mixed morphology and unexpected mtDNA lineages for their habitat (Garcia-Rodriguez et al., 1998; Vianna et al., 2006).

Out of the eight samples, three manatees from FG were morphologically classified as *T. manatus* (presence of nails, absence of white patch on the chest), but presented mtDNA haplotypes typically found in *T. inunguis* (haplotypes T2 and R1). Furthermore, microsatellite data confirmed the hybrid status of four of the eight manatees (two from FG and two from Brazil), and a karyotype of a single individual from Amapá (Brazil) indicated that it was a likely second-generation hybrid, a male offspring between a female F1 hybrid and a male *T. manatus* (Vianna et al., 2006). Due to the supposedly high prevalence of hybrids along the Guianas coast, this region located west of the Amazon River mouth was suggested to be a hybrid zone where *T. manatus* and *T. inunguis* individuals may contact and produce hybrids, further isolating the two *T. manatus* ESUs (Santos et al., 2016).

Hybridization has been recognized as a widespread phenomenon within animals, and it plays an important role in species evolution (Mallet, 2005). In endangered species, hybridization is often a consequence from declining population numbers due to anthropogenic activities (Fitzpatrick et al., 2015). When populations of closely related species – that have close or overlapping distribution and can still interbreed – decreases in size and consequently mating partners are rare, species hybridization is a likely outcome. Hybridization and introgression can have negative effects for a population (e.g. outbreeding depression), and the rise of hybrid individuals can further complicate the endangerment status of a species. If hybrids frequency is shown to be increased due to anthropogenic population decline of parental species, other alternatives should be discussed to avoid further decline of rare parental species and populations. However, in populations where hybridization is a natural phenomenon and does not pose a threat to the non-hybrid populations, improved management strategies can be developed to ensure the protection of hybrids. However, hybrids are usually not taken into consideration in conservation regulation and plans (Stronen & Paquet, 2013), and therefore the specimens and their habitat may be not eligible to be protected (but see Allendorf et al. 2001). Therefore, understanding hybridization events (i.e. their extension in space and time, ecological factors, interbreed, survival rate etc) and their causes (i.e. anthropogenic *versus* natural) is fundamental so that hybrids can be appreciated in conservation efforts.

In this study, we used a genomic approach to (i) investigate a previously recognized hybrid zone between manatee species, in order to (ii) understand the extent of hybridization, and (iii) to discuss ecological and evolutionary hypothesis for their occurrence and conservation implications.

## Methods

### Sampling

Tissue material from eight samples from French Guiana were collected from stranded animals, found dead on beaches and/or in mangroves. If the carcasses were not too degraded, phenotypical information was collected, such as presence of nails, skin color and extant of white patches on the chest and belly, expected to differentiate West Indian and Amazonian manatees. Kidney and liver samples were obtained from a dead *T. manatus* individual from the Tierpark Berlin Zoo, a born in captivity descendant of maternal grandparents from Georgetown, Guyana, and unknown father (Bell, 2001). Two *T. inunguis*, two *T. manatus* and a likely F2 hybrid from Amapá, Brazil (Vianna et al., 2006) were also included.

### DNA extraction and Sanger sequencing

DNA extractions were performed using the Dneasy Blood and Tissue kit (Qiagen). The mtDNA D-loop region was amplified using the primers L15926 and H16498 (Kocher et al., 1989). The amplification cycle followed Vianna et al. (2006)□, with annealing temperature of 55 °C. Nuclear genes (Apolipoprotein B - APOB, Amyloid precursor protein - APP, BMI1 proto-oncogene – BMI1, and CAMP Responsive element modulator – CREM), were amplified following Murphy et al. (2001) (Table S1). New primers for the APOB gene were designed using as reference the *T. manatus* GenBank sequence JN413954 (Meredith et al., 2011). For all loci, the PCR mix was prepared with final volume of 25 μL, containing 1X Taq reaction buffer; 1.5 mM of MgCl_2_; 200 μM of dNTPs; 0.5 μM of each primer; 0.5 U of Platinum Taq DNA Polymerase (Thermo Fisher Scientific). PCR products were purified by the polyethylene glycol (PEG) method (20% PEG 8000, 2.5M NaCl) (Sambrook, Fritsch, & Maniatis, 1989) and sequenced on the ABI 3130xl Genetic Analysis (Applied Biosystems) using the BigDye Termination v3.1 Cycle Sequencing kit. The same primers used for amplification were used for DNA sequencing. Chromatograms were analyzed in SeqScape v2.6 and consensus sequences were aligned using the ClustalX (Larkin et al., 2007) algorithm in the software MEGA 6 (Tamura et al., 2013). The relationships between haplotypes were inferred with the software NETWORK (Fluxus-engineering) using the median-joining algorithm (Bandelt, Forster, & Röhl, 1999). Summary statistics were calculated in DNAsp v5 (Rozas et al., 2003).

### Double-digest RADseq

Double-digest RAD (ddRAD) libraries were prepared following a previous protocol (Peterson et al., 2012) with adapter modifications (Meyer & Kircher, 2010), including two *T. inunguis* from Brazil, five manatees from French Guiana, and a *T. manatus* from Tierpark Zoo (Germany). In brief, 1 μg of genomic DNA was digested with the enzymes *Mse*I and *Eco*RI at 37 °C for at least two hours in a reaction volume proportional of DNA. The ligation of adapters was performed immediately after the digestion. Inline barcodes of varying sizes (5 to 9 bp) were added to the P5 adapter. The adapters and overhangs matching the enzymes were added. A sixteen-cycle indexing PCR was then performed in which one of the 50 barcodes described by Meyer and Kircher (2010) was added to the P7 adapter, was then cleaned with 1.5X AMPure XP magnetic beads and quantify in the Qubit 2.0 using the dsDNA HS assay (Life Technologies) and checked in the Agilent 2100 Bioanalyzer High Sensitivity DNA (Agilent Technologies). The indexed samples were then equimolarly pooled in two libraries and selected by size in BluePippin using the 1.5% cassette with R2 marker (Sage Science). We select the sequences in a range between 350-400 bp and checked with the Agilent Bioanalyzer 2100 High Sensitivity DNA chip. No PCR was performed after the samples were pooled to avoid chimera formation (DaCosta & Sorenson, 2014). Then the libraries were characterized using the KAPA SYBR^®^ FAST (Kapa Biosystems) kit and run in the in-house Illumina NextSeq 500 using a 300-cycle mid output v2 kit or in the in-house MiSeq with a 300-cycle kit.

Illumina reads were first demultiplexed according to P7 indexes. The software ipyrad 0.7.19 (Eaton, 2014) was used to demultiplex the P5 inline barcodes, trim the adapters and quality trimming. Since samples from FG were in an advanced state of decomposition, we first estimated the contamination level in each sample. A subsample of 20,000 merged sequences was randomly extracted and blasted against the NCBI nt database. BLAST results were visualized in MEGAN6 (Huson et al., 2007) and levels of bacterial contamination were evaluated. Samples that remained in the analysis after evaluating the levels of contamination were submitted to a stepwise filtering process. First, all reads were blasted (blastn) against the Florida manatee genome (GenBank Accession Number GCA000243295) and to the African elephant genome (GCA000001905). Sequences with blast hits below an e-value threshold of 10e-50 were retained. Subsequently, reads with no hits to either genome was blasted against a *Clostridium noyi* genome (see Results for reasoning) with a similar threshold. The remaining reads were finally blasted against a database of 8344 bacterial genomes and all reads presenting hits below the threshold were excluded from the clustering step. Clustering and SNP calling were performed in ipyrad. We used the following parameters: minimum coverage for a cluster: 8; clustering threshold: 0.90; data type: pairddrad; minimum samples in a final locus: 6. The remaining parameters were left as default.

Supposedly hybrid individuals used for ddRAD were originally from the Guianas region where the hybrid zone was reported (Vianna et al., 2006). As controls we used two Amazon manatees (*T. inunguis*) from Brazil and a West Indian manatee (*T. manatus*) represented by a Florida manatee (*T. manatus latirostris*) genome (Foote et al., 2015). The consensus sequences of each RAD locus were mapped to the Florida manatee genome using bowtie2 (Langmead & Salzberg, 2012) in order to identify the homologous regions. In parallel, all raw reads available in GenBank were mapped back to the genome contig sequences, also using bowtie2. Resulting SAM files were analyzed using customized python scripts for the identification of alleles at the target loci. The Florida manatee genotypic information was then added to the ddRAD results for subsequent analyses.

The software STRUCTURE 2.3.4 (Pritchard, Stephens, & Donnelly, 2000) was used to identify hybrids and possible different genetic groups. A total of twenty replicates were performed for each value of K from 1 to 5 (burn-in period: 20,000, 2 × 10^6^ iterations) under the admixture model and the assumption of correlated allele frequencies among populations. The USEPOPINFO model was used to specify the two parental species (POPFLAG=1) by assigning the two *T. inunguis* samples from Brazil and the *T. manatus* from Florida, USA as learning samples, while ancestry was estimated for all FG samples. The software CLUMPAK (Kopelman et al., 2015) was used to summary and visualize the results. Samples identified as pure individuals were subsequently assigned as learning samples. The observed heterozygosity (Hobs) was estimated using the R package adegenet 2.1.0 (Jombart & Ahmed, 2011). The number of differences between samples were estimated in the software Arlequin 3.5 (Excoffier & Lischer, 2010).

Hybridization simulations, using species-specific SNPs between the two-parental species, were performed using the command *hybridize* in adegenet (Fig. S1).

## Results

### Sanger sequencing

A total of 410 bp of mtDNA D-loop and 1,919 bp of nuclear DNA were sequenced, although four samples (M348, M827, M2108 and M2772) were partially sequenced for some nuclear genes (Table 1). Alleles were considered as species-specific based on population studies using larger samples of both species (Vianna et al. 2006; Santos et al. 2016, and unpublished data). Summary statistics and diagnostic polymorphisms for all sequenced loci can be seen in Tables S2 and S3.

**Table 1.** Sampling location and main results obtained for each sample. MtDNA D-loop haplotypes and nuclear haplotypes belonging to each species (*T. manatus*/*T. inunguis*, respectively) are shown in parenthesis.

| Sample | Sampling location | D-loop | Nuclear | ddRAD |
| --- | --- | --- | --- | --- |
| M113 | Paraná do Castanho, AM, Brazil | *T. inunguis* (T) | *T. inunguis* (0/8) | *T. inunguis* |
| M117 | Tefé, AM, Brazil | *T. inunguis* (Q2) | *T. inunguis* (0/8) | *T. inunguis* |
| M22 JAG | Awala, French Guiana | *T. manatus* (J2) | Introgressed (7/1) | Introgressed |
| M297 JAG | Awala, French Guiana | *T. inunguis* (T2) | *T. manatus* (8/0) | Introgressed |
| M348 JAG | Awala, French Guiana | *T. inunguis* (T2) | - | - |
| M423 JAG | Mana, French Guiana | *T. manatus* (J2) | Introgressed (7/1) | Introgressed |
| M524 JAG | Kourou, French Guiana | *T. inunguis* (T2) | *T. manatus* (8/0) | Introgressed |
| M827 JAG | Riviere des Cascades, French Guiana | *T. inunguis* (T2) | *T. inunguis* (0/2) | *T. inunguis* |
| M2108 JAG | Marina de Dégrad des Cannes, French Guiana | *T. inunguis* (T2) | - | - |
| M2772 JAG | Kaw River, French Guiana | *T. inunguis* (T2) | *T. manatus* (6/0) | - |
| Tierpark | Tierpark zoo, Berlin, Germany | *T. manatus* (M) | Introgressed (7/1) | *T. manatus* |
| M035 | Oiapoque, AP, Brasil | *T. Inunguis* (T) | Introgressed (6/2) | - |
| M042 | Baía Formosa, RN, Brazil | *T. manatus* (M) | *T. manatus* (8/0) | - |
| M047 | Beberibe, CE, Brazil | *T. manatus* (M) | *T. manatus* (8/0) | - |

*Trichechus manatus* and *T. inunguis s*amples from Brazil had both the mitochondrial and nuclear DNA consistent with their morphological classification and habitat, except for sample M035 (Table 1) from Amapá state (close to the frontier with French Guiana), which is a likely second-generation hybrid previously characterized by mtDNA, microsatellite and karyotype data (Santos et al., 2016). Most of the samples from FG had mixed nuclear and mtDNA from both species (Table 1). Surprisingly, a new D-loop mtDNA haplotype (J2), related to the *T. manatus* cluster II (Fig. S2), was found in two FG samples (M22 and M423). All five FG samples with *T. inunguis* mtDNA had the haplotype T2, previously described in FG samples as hybrids (Vianna et al., 2006). Remarkably, a FG manatee (M827) found in a tributary (Riviere des Cascades, 40 km from the river mouth at sea) of the Cayenne River had both *T. inunguis* mtDNA and nuclear genes (Table 1), which suggests a non-hybrid individual. The analysis of nuclear genes, also indicates that the *T. manatus* from Tierpark Zoo, which is maternally descendant from manatees from Suriname (in the Guianas coastline), presents 1 of 8 haplotypes derived from *T. inunguis*, thus a likely later generation hybrid born in captivity.

Most FG manatees were found dead in an advanced state of decomposition, but phenotypic information was obtained for two individuals. The individual M423 had a large white patch on its breast (Fig. S3) and no nails on its flippers, both characteristic of *T. inunguis*. The sample M524 had no nails but did not exhibit a white patch. Thus, both individuals presented phenotypes indicative of *T. inunguis* ancestry.

### ddRAD

A total of 30,486,215 paired sequences were obtained for eleven *Trichechus* samples. Eight samples presented similarities with the bacterial genomes (<1%), indicating a negligible contamination by decomposer organisms. On the other hand, three samples (M2772, M2108, M348) were mostly composed of bacterial sequences, primarily from the genus *Clostridium* and were excluded from further analysis. One sample (M827) had a high presence of bacterial reads from the genus *Clostridium* (Fig. S4), but due to the large number of reads, we were able to filter out bacterial sequences and include this sample in further analysis. Due to the elevated number of reads compared to other samples, a subsample of 2 million reads was used for sample M524.

After filtering for bacterial reads, samples had an average of 1,379,455 paired reads, although three samples had significant numbers of small fragment carry-over (DaCosta & Sorenson, 2014). All samples had overlapping fragments between 210-260 bp, and therefore all analyses were conducted within this range. Coverage depth varied greatly between clusters (Table S4), with many loci sequenced at low depth. Despite passing the paralog filter in ipyrad, five loci blasted in two separate regions of the *T. m. latirostris* genome and were excluded. The final dataset consisted of 947 clusters and 2287 SNPs. Genotypic data from the *T. m. latirostris* genome was obtained for 900 RAD loci and a total of 118 SNPs was unique to this.

A total of 720 unlinked SNPs was used in the STRUCTURE analysis. The preferred K was 2 (Fig. 1A). The results are in accordance with the four sequenced autosomal loci, the FG sample M827 is a non-hybrid *T. inunguis*, the sample from Tierpark is likely a non-hybrid (or highly-introgressed) *T. manatus* and all samples FG samples showed admixture between the two parental species (Fig. 1A and 1B). Furthermore, the assignment of M827 and Tierpark as training samples (added separately or combined) did not change the results in respect to hybrid admixture proportions (Fig. 1C).

**Figure 1.**
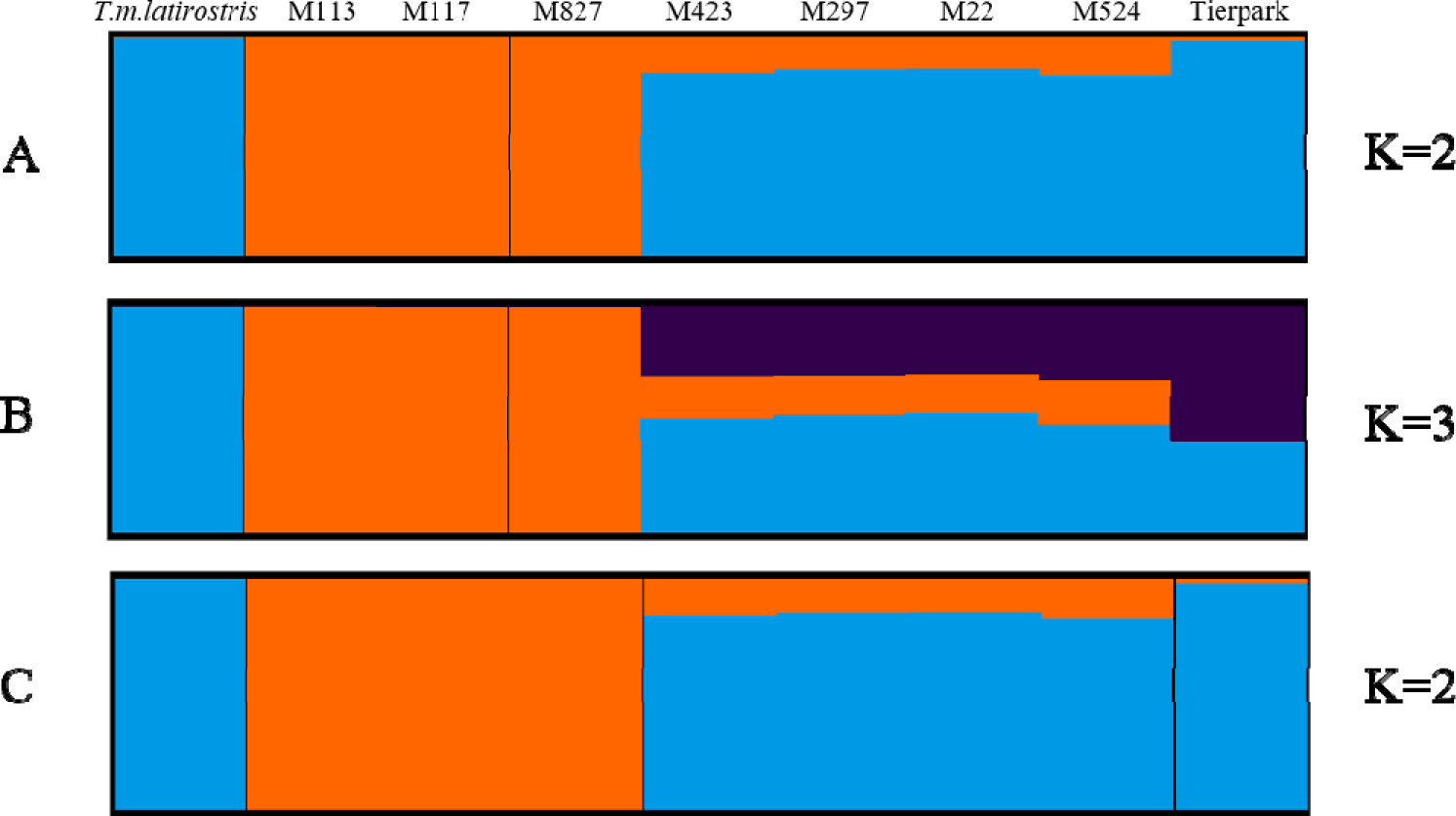
STRUCTURE results for unlinked SNPs. A and B, Admixture results using *T. m. latirostris* and *T. inunguis* from Brazil (M113 and M117) as training samples. C, Admixture results adding M827 and Tierpark as training samples.

The differences within samples (Table 2 and Table S5) showed a lower diversity in *T. inunguis* and *T. manatus* (Florida) when compared to *T. manatus* (Tierpark Zoo). Hybrid samples from FG were more similar to *T. manatus* from Tierpark (with maternal ancestors from Suriname, Guianas coastline) than from Florida (*T. m. latirostris*). The newly identified *T. inunguis* from FG (M827) was more closely related to *T. inunguis* from Brazil than to other FG samples, although it presents the typical mtDNA haplotype found in FG hybrids (Table 1).

**Table 2.** Pairwise differences between manatee samples considering the 720 unlinked SNPs. Both *T. inunguis* samples from Brazil were considered as one group. Upper diagonal: Corrected pairwise differences between samples; Diagonal: Number of polymorphic sites within samples; Lower diagonal: Pairwise divergence between samples.

|  | M827 | M423 | M297 | M22 | M524 | *T. manatus*<br>(Tierpark) | *T. inunguis* | *T. manatus*<br>(Florida) |
| --- | --- | --- | --- | --- | --- | --- | --- | --- |
| M827 | 24 | 73 | 77 | 74 | 83 | 86 | 0 | 134 |
| M423 | 0.62 | 64 | 0 | 0 | 0 | 13 | 101 | 35 |
| M297 | 0.67 | 0.00 | 52 | 0 | 1 | 17 | 102 | 36 |
| M22 | 0.61 | 0.00 | 0.00 | 70 | 0 | 10 | 95 | 29 |
| M524 | 0.61 | 0.00 | 0.01 | 0.00 | 84 | 8 | 109 | 37 |
| *T. manatus*<br>(Tierpark) | 0.70 | 0.19 | 0.25 | 0.14 | 0.10 | 49 | 108 | 41 |
| *T. inunguis* | 0.00 | 0.77 | 0.79 | 0.75 | 0.76 | 0.80 | 21 | 157 |
| *T. manatus*<br>(Florida) | 0.90 | 0.50 | 0.55 | 0.43 | 0.45 | 0.60 | 0.90 | 6 |

The simulations of hybridization scenarios (Fig. S1) and the STRUCTURE results showed a similar level of admixture between hybrids individuals (>F2).

## Discussion

This paper explores the genetic diversity of manatees at the frontier region of the distribution areas of two species, along the Guianas coastline with some characteristics of a large estuarine environment close to the Amazon River mouth. Here we (i) validate a pipeline to extract genotypic information from the raw reads of full genomes, overcoming samples limitations for ddRAD analysis, (ii) describe an incongruence between mitochondrial and nuclear DNA in FG manatees, supporting the extent of the hybridization in this population, (iii) report the first *T. inunguis* outside the Amazon Basin, in a French Guiana river at least 40 km away from the sea and (iii) discuss the implications for conservation and management of this hybrid population under influence of the Amazon River plume.

### Low-quality DNA sample quality for population genomics

The combination of high-coverage sequencing and bioinformatic filtering allowed the use of low-quality samples from decomposing animals found in FG rivers. The high number of bacterial reads was an indicative of the low sample quality. The presence of *Clostridium, Lactobacillus* and other Enterobacteriaceae sequences was previously associated with decomposing bodies (Hyde et al., 2013). The samples composed mostly of bacterial reads showed a higher bacterial diversity than the other samples, which could indicate a more advanced of decomposition than the others. Nevertheless, ddRAD data can be used to assess sample quality, local bacterial diversity, and bacterial composition of decaying samples. The use of decomposing animals, can be the only biological material usable to assess the genetic of small populations or that have difficulties in sampling. In this study, they have proven to be a valuable tool to identify SNPs to investigate the hybrid swarming phenomenon occurring along the Guianas coastline (Santos et al., 2016).

### Manatees in French Guiana: a hybrid population

The presence of a *T. manatus* mtDNA clade II haplotype in FG, combined with the presence of *T. inunguis* individuals, is an indication that this population has a more complex dynamic than previously thought. Previous studies have identified that the distribution limit between *T. manatus* clades II and III was between Venezuela and Guyana (Vianna et al., 2006), where a possible historical barrier to gene flow was hypothesized. The presence of clade II in FG highlights the gene flow of western manatees to the hybrid zone, where they meet eastern manatees (from Brazil), as shown by the presence of clade III mtDNA lineages (L, M, N, O) in Guyana (Garcia-Rodriguez et al., 1998), the westernmost country of the Guianas coastline. Thus, this *T. manatus vs T. inunguis* hybrid zone is also a secondary contact zone between both *T. manatus* ESUs (Vianna et al., 2006; Santos et al., 2016).

The extent of the hybrid zone between *T. manatus* and *T. inunguis* as suggested earlier (Vianna et al., 2006; Santos et al., 2016), and the unusual high rate of hybrids supports a possible ecological and biological advantage of hybrids in this peculiar environment. The identification of later generation hybrid manatees indicates that this region may have an old history of hybridization, and hybrids may present higher fitness in the local ecological conditions. This remarkable finding of an extended distribution of a large aquatic mammal throughout riverine systems separated by hundreds of kilometers of sea implies the unique influence of the largest river basin in the world. The Amazon River discharge at the Atlantic Ocean shapes the distribution of many taxa of the marine biota (Spalding et al., 2007), sometimes promoting vicariance, as it can be seen for the disjoint distribution of West Indian manatees (Santos et al., 2016; Barros et al., 2017), with eastern and western ESUs separated by the hybrid zone acting as a barrier to gene flow. However, the presence of the Amazon River plume seems also to promote the connection of freshwater biota from separate river basins, like catfishes (Barthem et al., 2017), as we have also observed for the Amazonia manatee found in a river in French Guiana.

### The Amazonian manatee outside the Amazon basin

The presence of *T. inunguis* reported here for the first time outside the Amazon basin indicates a possible connection between freshwater systems along the Guianas coastline and the Amazon River, in a region under influence of the Amazon plume that runs westwards because of the North Brazil Current (Muller-Karger, McClain, & Richardson, 1988). Indeed, the identification of a large Amazon River mammal in a French Guiana freshwater system supports the idea of a broad and unique low-salinity estuarine-like habitat along the Guianas coastline that connects freshwater biodiversity of further apart river basins under the influence of the Amazon River plume (Muller-Karger, McClain, & Richardson, 1988; Artigas et al., 2003; Anthony et al., 2013; Gensac et al., 2016; Barthem et al., 2017). From the Amazon River mouth to the west of the Guianas coast, the suitable habitats for *T. manatus* are narrow, due to the quite restricted alluvial coastal plain. The fluvial sediment supplies of the Amazon River shaped the coastal landscape during the Quaternary (Anthony et al., 2013; Gensac et al., 2016) and resulted in a continuous mangrove-dominated ecosystem along the Guianas coastline from Amapá (Brazil) to Suriname, with rather close salinity and turbidity traits brought by the Amazon plume. This coastal geomorphology associated with the estuary-like characteristics may thus provide an ecological continuum (Artigas et al., 2003), from the currently accepted distribution of *T. inunguis* in the Amazon Basin towards the area of occurrence of *T. manatus* along the Guianas coastline. Furthermore, the North Brazil Current and the Amazon River plume (Froidefond et al., 2002; Fratantoni & Richardson, 2006) likely promote a westward movement of animals from the Amazon River mouth.

The newly reported *T. inunguis* in this study has the same mtDNA haplotype as the hybrids and has only been described in FG (Vianna et al., 2006), which implies this hybridization may be a local phenomenon, involving both species. The mtDNA haplotypes and the observed heterozygosity from simulated hybrids indicates that the hybridization phenomenon has been occurring for several generations and that later-generation hybrids in FG tend to mate with each other, thus initial hybrids (F1 and F2) are at least partially fertile, particularly the females, as occurs in other mammals (Allendorf et al., 2001; Stronen & Paquet, 2013; Fitzpatrick et al., 2015). Future sampling efforts at larger geographic extent could help understanding the ratio of hybrids to “pure” manatees in the Guianas coastline, and if hybrids from different generations (F1, F2, etc) are also present in the population.

### Implications for conservation

The need for conservation of hybrids is a controversial and challenging issue. Within species, Conservation Units are independent population units that are used to help guide management and conservation initiative, at the more relevant spatial extent (Funk et al., 2012). Other commonly discussed conservation units correspond to evolutionarily significant units (ESUs) and management units (MUs). An ESU can be defined as a population exhibiting high genetic and ecological distinctiveness (Crandall et al., 2000) as suggested for *T. manatus* (Vianna et al., 2006; Santos et al., 2016), while MUs are expected to be demographically independent (Palsbøll, Bérubé, & Allendorf, 2007).

Distinctive genetic signatures of the Guianan manatees and their likely adaptive characters to the ecological conditions of this peculiar coastline and drainages may allow considering them as a special Conservation Unit, as a distinct MU, if not even an ESU. In this case, this population would require a particular conservation effort. Even though global IUCN do not consider hybrids as relevant taxa to be evaluated (IUCN Standards and Petitions Subcommittee, 2017), regional Red List assessment in French Guiana (UICN France et al., 2017) concluded it to be in a stable or decreasing trend, with a small population size due to restricted habitats, and ongoing threats such as incidental poaching, and to be included in an “endangered” conservation status.

Thus, our study shows that French Guiana (and from other neighboring Guianan coasts) manatees are hybrids between *Trichechus manatus*, the coastal (and associated freshwater environment) species and *T. inunguis*, the freshwater adapted species from the Amazon River basin. Because genomic contribution to the local hybrids mostly comes from *T. manatus*, we conclude this hybridization is occurring for many generations with the input of *T. inunguis* individuals surviving along the Amazon River plume. The extent of the hybrid zone within the distribution area of *T. manatus* raises the question of the likely ecological advantages of this introgression of genes from a freshwater adapted genome. The understanding of associated genetic process cause could be investigated by searching for natural selection signs throughout the genome of both species and hybrids, which could also give clues for the putative dominance of characters and directionality in interbreeding.

If this particular aquatic ecosystem promoted by the Amazon plume indeed influences the distribution and population dynamics of many other species along this coastal area, a regional conservation program should be devised, with an emphasis to preserve this unique aquatic environment of the Guiana shield coastline

## Acknowledgments

We thank Susan Mbedi (BeGenDiv) and Hannah Ebbinghaus for their technical support in next-generation sequencing and Max Driller for bioinformatic support. Claudia Szentkis (IZW) for providing the sample of the captive manatee. STV was supported by an Alexander von Humboldt Foundation fellowship, CSL by a CAPES (Brazil) fellowship, and FRS by a CNPq (Brazil) research fellowship. This project was performed under ICMBio/SISBIO permit 45028 and received funding support from FAPEMIG, CNPq and Fundação o Boticário from Brazil. Samples from Brazil are kept in the collection of the Laboratório de Biodiversidade e Evolução Molecular in UFMG, Belo Horizonte, with a collaboration with ICMBio and Instituto de Desenvolvimento Sustentável Mamirauá, as part of the National Action Plan for Conservation of Sirenians of Brazil. The use of genetic resources for this study is declared in the SisGen platform according to Brazilian law 13123/2015.

Samples from French Guiana are kept in the databank JAGUARS, at Cayenne, French Guiana. The JAGUARS tissue collection is supported by the INDIGEN project, funded by European Community (ERDF funds), the Collectivité Territoriale de Guyane, and the DEAL Guyane.

The use of the genetic resources was declared to the French Ministry of Environment under the reference TSP 48704, in compliance with the Access and Benefit Sharing procedure implemented by the Loi pour la Reconquête de la Biodiversité.

## References

Allendorf, F. W., Leary, R. F., Spruell, P., & Wenburg, J. K. (2001). The problems with hybrids: setting conservation guidelines 16, 613–622.

Anthony, E. J., Gardel, A., Proisy, C., Fromard, F., Gensac, E., Peron, C., Walcker, R., & Lesourd, S. (2013). The role of fluvial sediment supply and river-mouth hydrology in the dynamics of the muddy, Amazon-dominated Amapá-Guianas coast, South America: A three-point research agenda. J. South Am. Earth Sci. 44, 18–24.

Artigas, L. F., Vendeville, P., Leopold, M., Guiral, D., & Ternon, J.-F. (2003). Marine biodiversity in French Guiana: Estuarine, coastal, and shelf ecosystems under de influence of Amazonian waters. Gayana (Concepción) 67, 302–326.

Bandelt, H. J., Forster, P., & Röhl, A. (1999). Median-joining networks for inferring intraspecific phylogenies. Mol. Biol. Evol. 16, 37–48.

Barros, H. M. D. d. R., Meirelles, A. C. O., Luna, F. O., Marmontel, M., Cordeiro-Estrela, P., Santos, N., & Astúa, D. (2017). Cranial and chromosomal geographic variation in manatees (Mammalia: Sirenia: Trichechidae) with the description of the Antillean manatee karyotype in Brazil. J. Zool. Syst. Evol. Res. 55, 73–87.

Barthem, R. B., Goulding, M., Leite, R. G., Cañas, C., Forsberg, B., Venticinque, E., Petry, P., Ribeiro, M. L. D. B., Chuctaya, J., & Mercado, A. (2017). Goliath catfish spawning in the far western Amazon confirmed by the distribution of mature adults, drifting larvae and migrating juveniles. Sci. Rep. 7, 1–13.

Bell, C. E. (2001). Encyclopedia of the World’s Zoos. Vol. 1. Taylor & Francis.

Castelblanco-Martínez, D. N., dos Reis, V., & de Thoisy, B. (2017). How to detect an elusive aquatic mammal in complex environments? A study of the Endangered Antillean manatee Trichechus manatus manatus in French Guiana. Oryx 1–11.

Castelblanco-Martinez, D. N., Rosas, F. C. W., Bermudez, A., & Trujillo-Gonzales, T. (2003). Conservation status of the West Indian manatee, Trichechus manatus manatus, in the Middle Orinoco (Vichada, Colombia). In Conf. Proceedings, 15th Bienn. Conf. Biol. Mar. Mammals, Greensboro, North Carolina. p. 30.

Crandall, K. A., Bininda-emonds, O. R. P., Mace, G. M., & Wayne, R. K. (2000). Considering evolutionary processes in Conservation Biology. Trends Ecol. Evol. 15, 290–295.

DaCosta, J. M., & Sorenson, M. D. (2014). Amplification Biases and Consistent Recovery of Loci in a Double-Digest RAD-seq Protocol. PLoS One 9, 1–14.

de Thoisy, B., Spiegelberger, T., Rousseau, S., Talvy, G., Vogel, I., & Vié, J.-C. (2003). Distribution, habitat, and conservation status of the West Indian manatee Trichechus manatus in French Guiana. Oryx 37, 431–436.

Deutsch, C.., Self-Sullivan, C., & Mignucci-Giannoni, A. (2008). The West Indian manatee, Trichechus manatus.

Eaton, D. A. R. (2014). PyRAD: Assembly of de novo RADseq loci for phylogenetic analyses. Bioinformatics 30, 1844–1849.

Excoffier, L., & Lischer, H. E. L. (2010). Arlequin suite ver 3.5: A new series of programs to perform population genetics analyses under Linux and Windows. Mol. Ecol. Resour. 10, 564–567.

Fitzpatrick, B. M., Johnson, J. R., Carter, E. T., & Arter, E. T. C. (2015). Hybridization and the species problem in conservation Hybridization and the species problem in conservation 61, 206–216.

Foote, A. D., Liu, Y., Thomas, G. W. C., Vinař, T., Alföldi, J., Deng, J., Dugan, S., van Elk, C. E., Hunter, M. E., Joshi, V., Khan, Z., Kovar, C., Lee, S. L., Lindblad-Toh, K., Mancia, A., Nielsen, R., Qin, X., Qu, J., Raney, B. J., Vijay, N., Wolf, J. B. W., Hahn, M. W., Muzny, D. M., Worley, K. C., Gilbert, M. T. P., & Gibbs, R. A. (2015). Convergent evolution of the genomes of marine mammals. Nat. Genet. 47, 272–275.

Fratantoni, D. M., & Richardson, P. L. (2006). The Evolution and Demise of North Brazil Current Rings*. J. Phys. Oceanogr. 36, 1241–1264.

Froidefond, J. M., Gardel, L., Guiral, D., Parra, M., & Ternon, J. F. (2002). Spectral remote sensing reflectances of coastal waters in French Guiana under the Amazon influence. Remote Sens. Environ. 80, 225–232.

Funk, C. W., McKay, J. K., Hohenlohe, P. A., & Allendorf, F. W. (2012). Harnessing genomics for delineating conservation units 27, 489–496.

Garcia-Rodriguez, a. I., Bowen, B. W., Domning, D., Mignucci-Giannoni, a. a., Marmontel, M., Montoya-Ospina, R. a., Morales-Vela, B., Rudin, M., Bonde, R. K., & McGUIRE, P. M. (1998). Phylogeography of the West Indian manatee (Trichechus manatus): how many populations and how many taxa? Mol. Ecol. 7, 1137–1149.

Gensac, E., Martinez, J. M., Vantrepotte, V., & Anthony, E. J. (2016). Seasonal and inter-annual dynamics of suspended sediment at the mouth of the Amazon river: The role of continental and oceanic forcing, and implications for coastal geomorphology and mud bank formation. Cont. Shelf Res. 118, 49–62.

Huson, D. H., Auch, A. F., Qi, J., & Schuster, S. C. (2007). MEGAN analysis of metagenomic data. Genome Res. 17, 377–386.

Hyde, E. R., Haarmann, D. P., Lynne, A. M., Bucheli, S. R., & Petrosino, J. F. (2013). The Living Dead: Bacterial Community Structure of a Cadaver at the Onset and End of the Bloat Stage of Decomposition. PLoS One 8, 1–10.

IUCN Standards and Petitions Subcommittee. (2017). Guidelines for Using the IUCN Red List Categories and Criteria. Version 13. Cambridge IUCN.

Jombart, T., & Ahmed, I. (2011). adegenet 1.3-1: New tools for the analysis of genome-wide SNP data. Bioinformatics 27, 3070–3071.

Kocher, T. D., Thomas, W. K., Meyer, A., Edwards, S. V., Paabo, S., Villablanca, F. X., & Wilson, A. C. (1989). Dynamics of mitochondrial DNA evolution in animals: amplification and sequencing with conserved primers. Proc. Natl. Acad. Sci. 86, 6196–6200.

Kopelman, N. M., Mayzel, J., Jakobsson, M., Rosenberg, N. A., & Mayrose, I. (2015). Clumpak: A program for identifying clustering modes and packaging population structure inferences across K. Mol. Ecol. Resour. 15, 1179–1191.

Langmead, B., & Salzberg, S. L. (2012). Fast gapped-read alignment with Bowtie 2. Nat Methods 9, 357–359.

Larkin, M., Blackshields, G., Brown, N., Chenna, R., McGettigan, P., McWilliam, H., Valentin, F., Wallace, I., Wilm, A., Lopez, R., Thompson, J., Gibson, T., & Higgins, D. (2007). ClustalW and ClustalX version 2. Bioinformatics 23, 2947–2948.

Mallet, J. (2005). Hybridization as an invasion of the genome. Trends Ecol. Evol. 20, 229–237.

Meredith, R. W., Janečka, J. E., Gatesy, J., Ryder, O. a, Fisher, C. a, Teeling, E. C., Goodbla, A., Eizirik, E., Simão, T. L. L., Stadler, T., Rabosky, D. L., Honeycutt, R. L., Flynn, J. J., Ingram, C. M., Steiner, C., Williams, T. L., Robinson, T. J., Burk-Herrick, A., Westerman, M., Ayoub, N. a, Springer, M. S., & Murphy, W. J. (2011). Impacts of the Cretaceous Terrestrial Revolution and KPg extinction on mammal diversification. Science (80-.). 334, 521–524.

Meyer, M., & Kircher, M. (2010). Illumina sequencing library preparation for highly multiplexed target capture and sequencing. Cold Spring Harb. Protoc. 1–7.

Montoya-ospina, R. A., Caicedo-herrera, D., Milla, S. L., Mignucci-giannoni, A. A., & Lefebvre, L. W. (2001). Status and distribution of the West Indian manatee, Trichechus manatus manatus, in Colombia 102, 117–129.

Muller-Karger, F. E., McClain, C. R., & Richardson, P. L. (1988). The dispersal of the Amazon’s water. Nature 332, 56–59.

Murphy, W. J., Eizirik, E., Johnson, W. E., Zhang, Y. P., Ryder, O. A., & O’Brien, S. J. (2001). Molecular phylogenetics and the origins of placental mammals. Nature 409, 614–618.

Palsbøll, P. J., Bérubé, M., & Allendorf, F. W. (2007). Identification of management units using population genetic data. Trends Ecol. Evol. 22, 11–16.

Peterson, B. K., Weber, J. N., Kay, E. H., Fisher, H. S., & Hoekstra, H. E. (2012). Double digest RADseq: An inexpensive method for de novo SNP discovery and genotyping in model and non-model species. PLoS One 7, 1–11.

Pritchard, J. K., Stephens, M., & Donnelly, P. (2000). Inference of population structure using multilocus genotype data. Genetics 155, 945–959.

Rozas, J., Sánchez-DelBarrio, J. C., Messeguer, X., & Rozas, R. (2003). DnaSP, DNA polymorphism analyses by the coalescent and other methods. Bioinformatics 19, 2496–2497.

Sambrook, J., Fritsch, E. F., & Maniatis, T. (1989). Molecular cloning. Society 68, 1232–1239.

Santos, F. R., Barros, H. M., Schetino, M. A., & Lima, C. S. (2016). Genetics. In A. C. Meirelles & V. L. Carvalho (Eds.), West Indian Manatee Biol. Conserv. Brazil / Peixe-boi Mar. Biol. e Conserv. no Bras. 1st ed., pp. 63–75. São Paulo, SP: Bambu Editora e Artes Gráfiacas.

Spalding, M. D., Fox, H. E., Allen, G. R., Davidson, N., Ferdaña, Z. A., Finlayson, M., Halpern, B. S., Jorge, M. A., Lombana, A., Lourie, S. A., Martin, K. D., Mcmanus, E., Molnar, J., Recchia, C. A., & Robertson, J. (2007). Marine Ecoregions of the World: A Bioregionalization of Coastal and Shelf Areas. Bioscience 57, 573–583.

Stronen, A. V., & Paquet, P. C. (2013). Perspectives on the conservation of wild hybrids. Biol. Conserv. 167, 390–395.

Tamura, K., Stecher, G., Peterson, D., Filipski, A., & Kumar, S. (2013). MEGA6: Molecular evolutionary genetics analysis version 6.0. Mol. Biol. Evol. 30, 2725–2729.

UICN France, MNHN, GEPOG, Kwata, Biotope, Hydreco, & OSL. (2017). La Liste rouge des espèces menacées en France - Chapitres de la Faune vertébrée de Guyane. Paris, France.

Vianna, J. A., Bonde, R. K., Caballero, S., Giraldo, J. P., Lima, R. P., Clark, A., Marmontel, M., Morales-Vela, B., De Souza, M. J., Parr, L., Rodríguez-Lopez, M. a, Mignucci-Giannoni, A. a, Powell, J. a, & Santos, F. R. (2006). Phylogeography, phylogeny and hybridization in trichechid sirenians: implications for manatee conservation. Mol. Ecol. 15, 433–447.

