## Supplementary Material for "Manatee genomics supports a special conservation area in the Guianas coastline under the influence of the Amazon River plume"

**Appendix**

**Table S1.** Primer sequences and annealing temperatures used for amplification and Sanger sequencing of nuclear and mitochondrial loci.

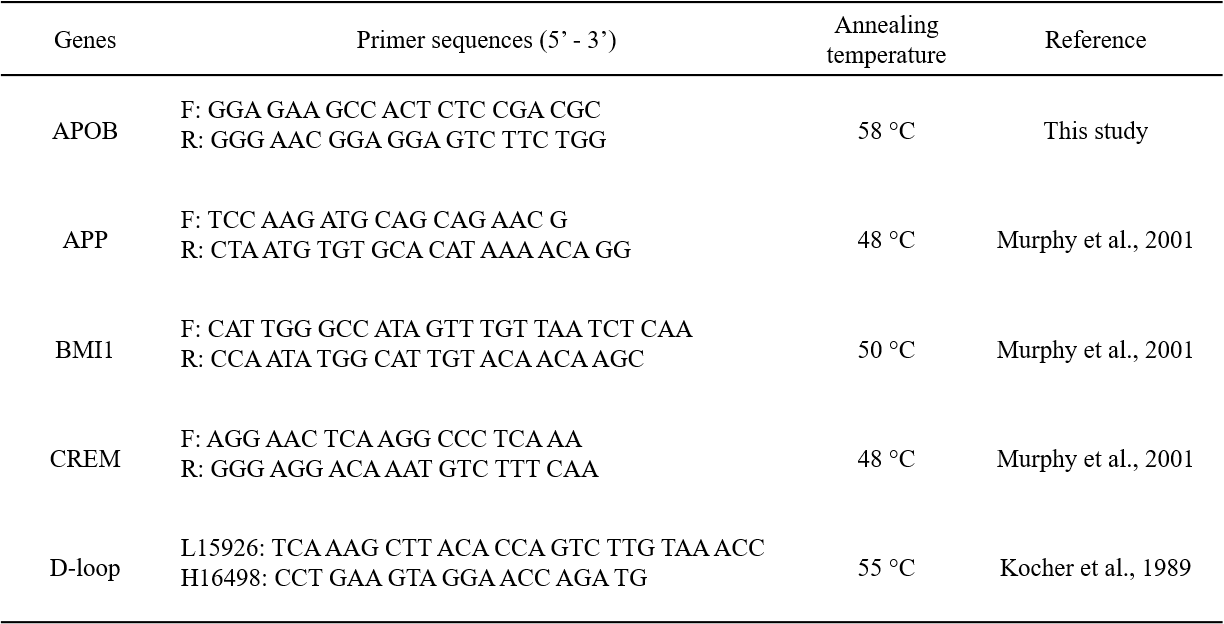

**Figure S1.** Simulated Observed heterozygosity for two different mating schemes. Black Circles represent results from the “Backcross” scenario, while black triangles represent crosses from the “Hybrid” scenario. The green square represent first generation hybrids. The red circle represent the observed heterozygosity of FG hybrids.

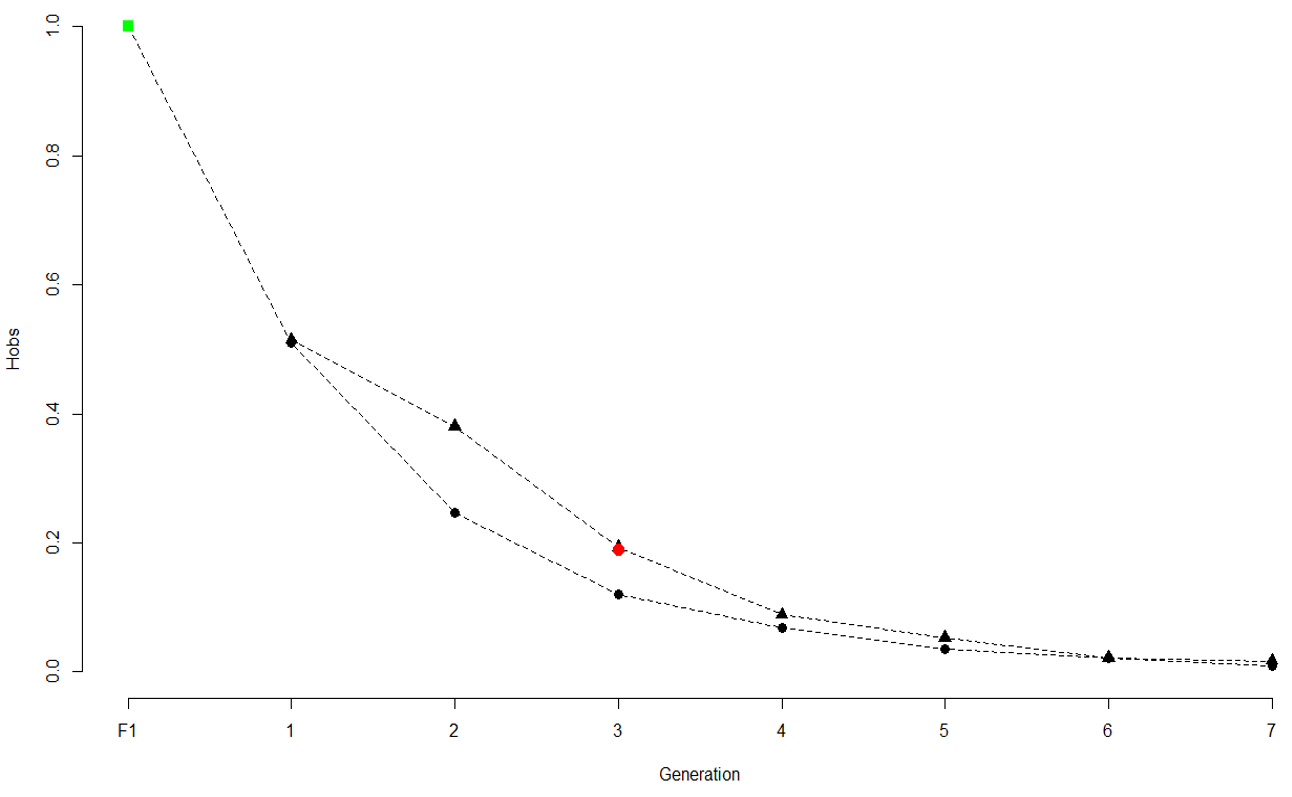

**Table S2.** Diagnostic sites for four nuclear Sanger sequenced loci. Polymorphisms restricted to *T. inunguis* are shown with an asterisk. Numbers correspond to polymorphic positions in each locus.

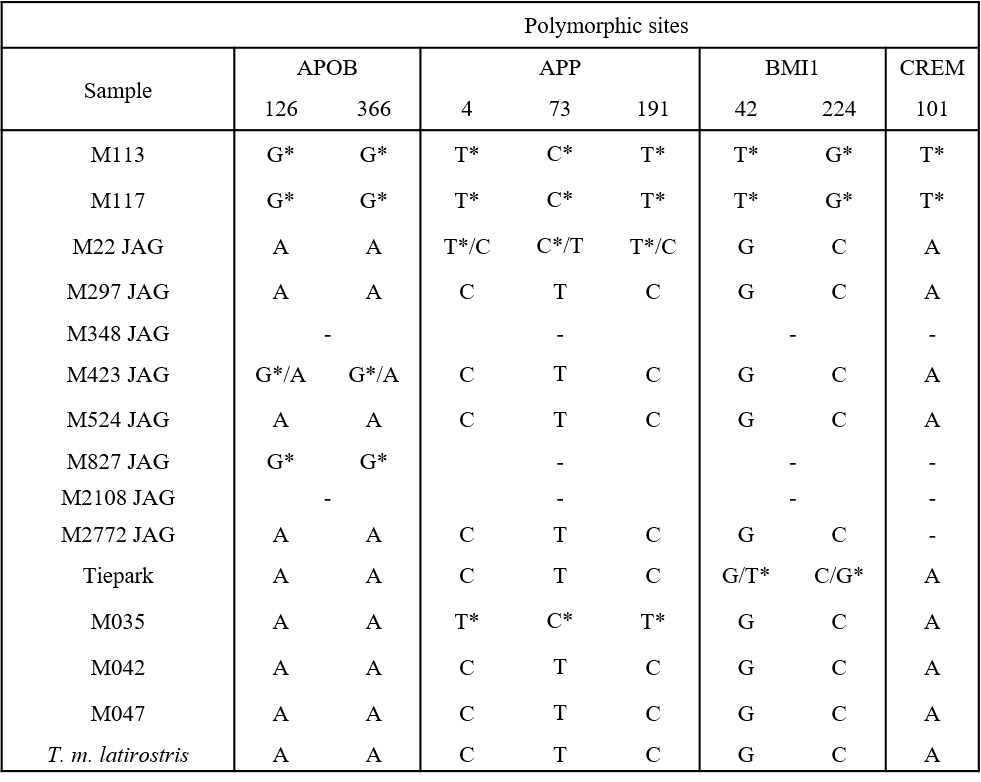

**Table S3.** Summary statistics for five Sanger sequenced loci. Only samples from French Guiana are included. Abbreviations: S, Number of variable sites; π, Nucleotide diversity; Hd, Haplotype diversity.

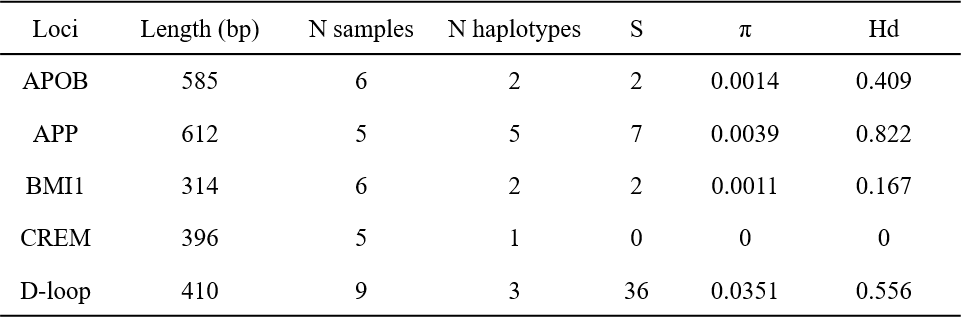

**Figure S2.** Median joining network tree showing the relationships between *T. manatus* D-loop haplotypes. Haplotype names follow Vianna et al. (2006). Number of mutations are indicated by numbers on branches. The new haplotype (J2) is shown in gray.

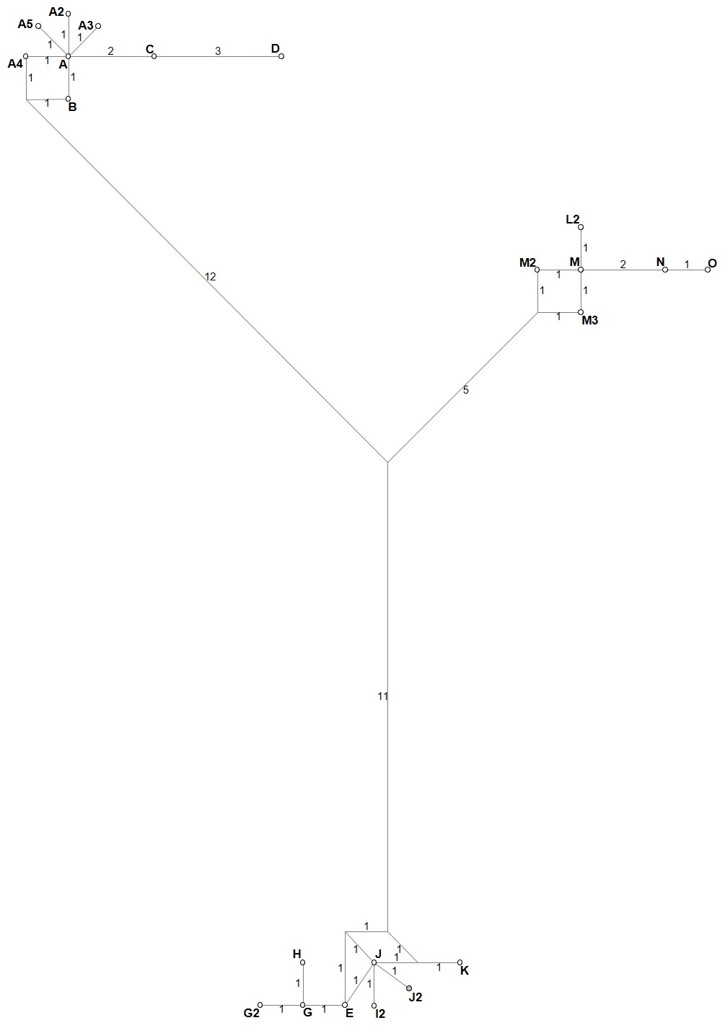

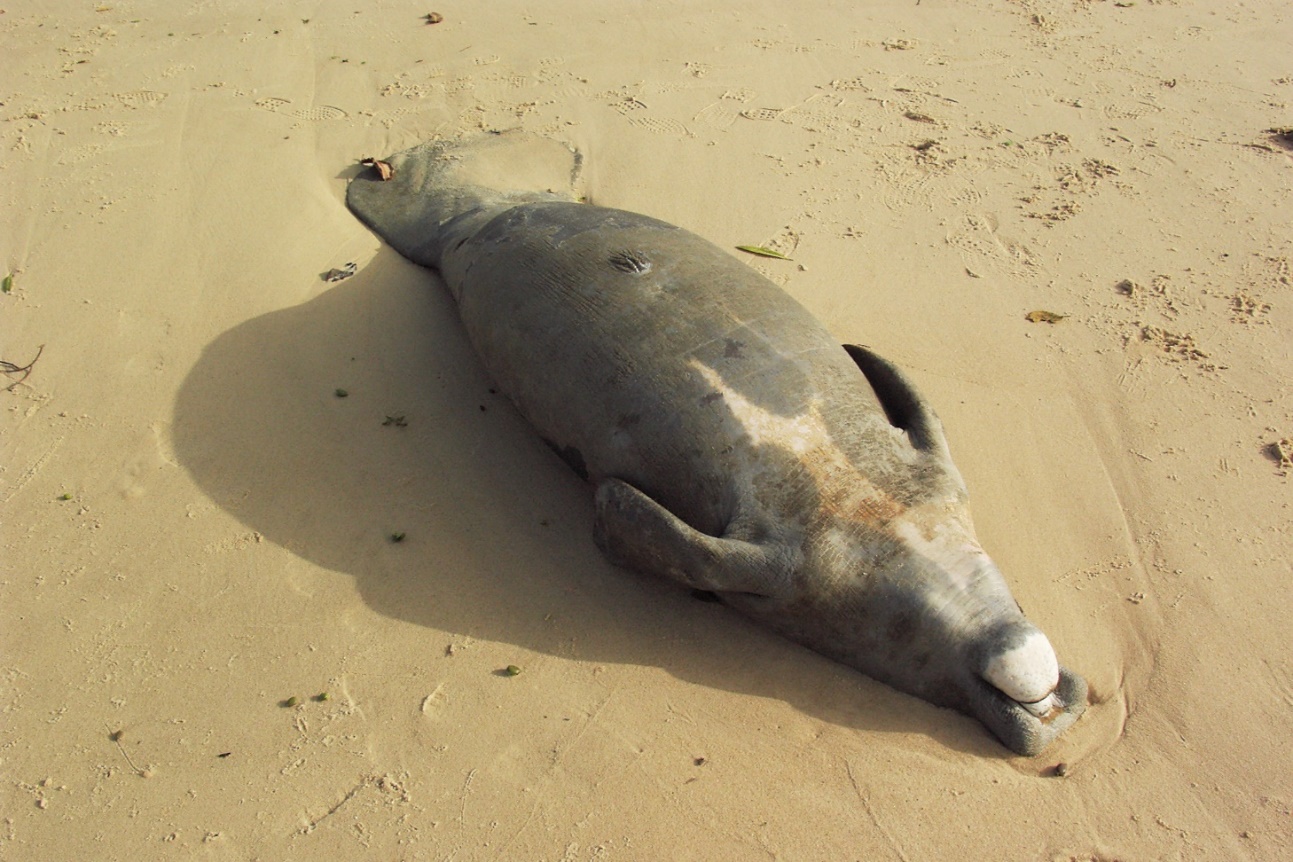
 **Figure S3.** Photo of the individual M423, collected in French Guiana, with a large white patch on its breast and no nails on its flippers.

**Figure S4.** Taxon composition from a random subsample of 20,000 reads for each sequenced manatee sample analysed in this study. Colors represent Blast results. Results were ranked according to Class.

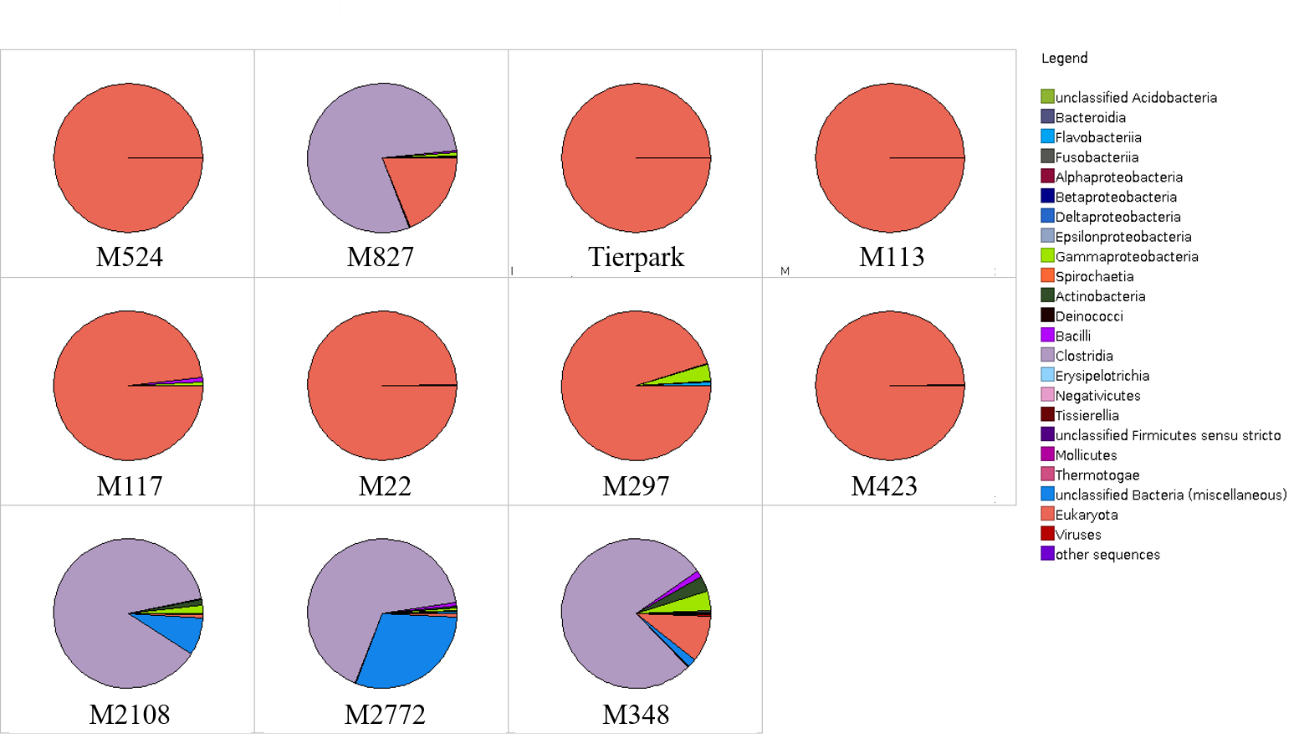

**Table S4.** Description of the ddRAD analysis. *N reads* represents the total number of reads obtained, *% bacterial reads* show the percentage of reads blasted against bacterial sequences out of a random subsample of 20,000 reads, *N reads used* show reads between 210-260 bp used for clustering after all filtering steps, N clusters is the total of clusters obtained per sample.

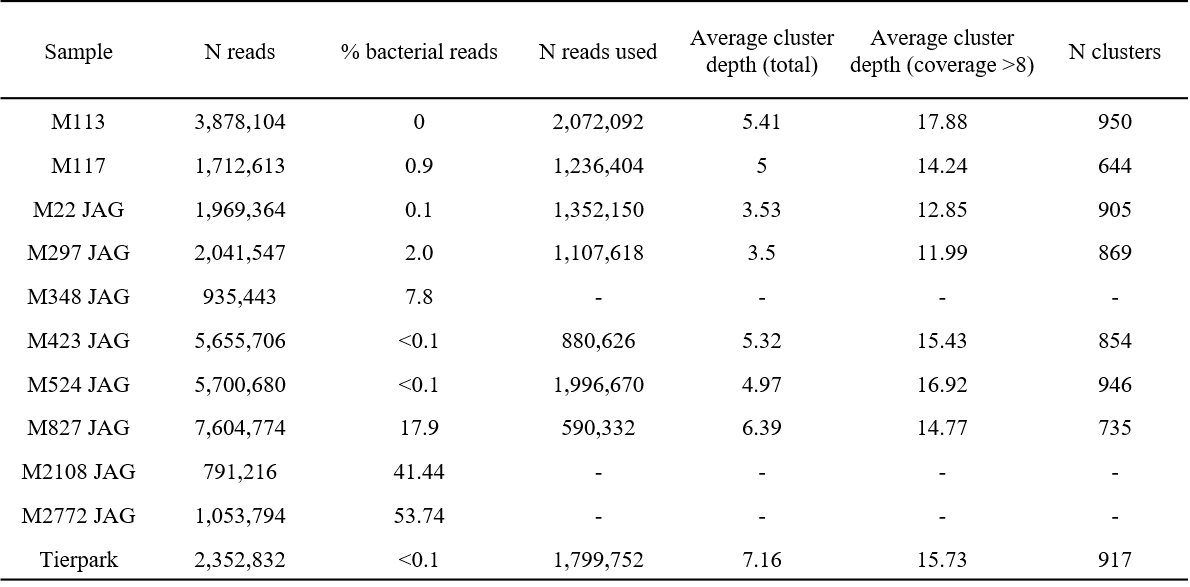

**Table S5.** Pairwise divergence between samples for the species-specific SNP dataset. Samples M113 and M117 were lumped into *T. inunguis*.

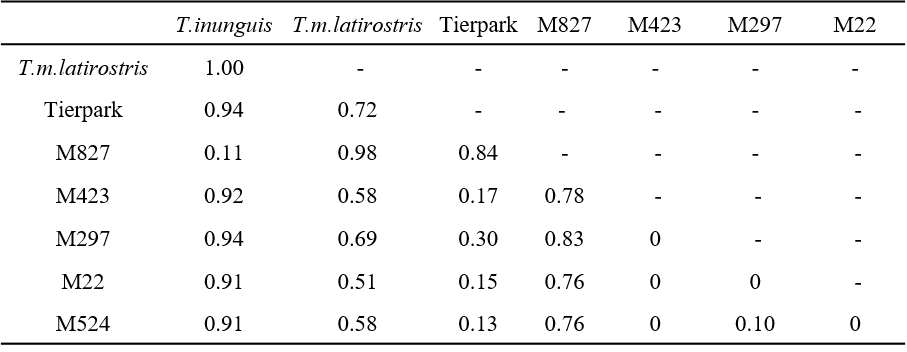
